# Predicting T cell activation based on intracellular calcium fluctuations

**DOI:** 10.1101/2023.06.14.545014

**Authors:** Sébastien This, Santiago Costantino, Heather J. Melichar

## Abstract

Adoptive T cell therapies rely on the transduction of T cells with a predetermined antigen receptor which redirects their specificity towards tumor-specific antigens. Despite the development of multiple platforms for tumor-specific T cell receptor (TCR) discovery, this process remains time consuming and skewed toward high-affinity TCRs. Specifically, the methods for identifying therapeutically-relevant TCR sequences, predominantly achieved through the enrichment of antigen-specific T cells, represents a major bottleneck for the broader application of TCR-engineered cell therapies. Fluctuation of intracellular calcium levels in T cells is a well described, proximal readout of TCR signaling. Hence, it is an attractive candidate marker for identifying antigen-specific T cells that does not require *in vitro* antigen-specific T cell expansion. However, calcium fluctuations downstream of TCR engagement with antigen are highly variable; we propose that appropriately-trained machine learning algorithms may allow for T cell classification from complex datasets such as those related to polyclonal T cell signaling events. Using deep learning tools, we demonstrate efficient and accurate prediction of antigen-specificity based on intracellular Ca^2+^ fluctuations of *in vitro*-stimulated CD8^+^ T cells. Using a simple co-culture assay to activate monoclonal TCR transgenic T cells of known specificity, we trained a convolutional neural network to predict T cell reactivity, and we test the algorithm against T cells bearing a distinct TCR transgene as well as a polyclonal T cell response. This approach provides the foundation for a new pipeline to fast-track antigen specific TCR sequence identification for use in adoptive T cell therapy.

**Significance Statement:** While T cells engineered to express a cancer-specific T cell receptor (TCR) are emerging as a viable approach for personalized therapies, the platforms for identifying clinically-relevant TCR sequences are often limited in the breadth of antigen receptors they identify or are cumbersome to implement on a personalized basis. Here, we show that imaging of intracellular calcium fluctuations downstream of TCR engagement with antigen can be used, in combination with artificial intelligence approaches, to accurately and efficiently predict T cell specificity. The development of cancer-specific T cell isolation methods based on early calcium fluctuations may avoid the biases of current methodologies for the isolation of patient-specific TCR sequences in the context of adoptive T cell therapy.

## Introduction

Adoptive T cell therapies are revolutionizing cancer treatment. In this context, T cells are engineered to redirect their specificity towards cancer antigens with either a chimeric antigen receptor or a predetermined, tumor-specific T cell receptor (TCR). TCR-T cell therapies have the potential to recognize antigens that arise from mutations, fusion proteins, and aberrantly expressed regions of the genome thereby increasing the breadth of targets (1, 2). However, the identification of tumor-specific TCRs is challenging due, in part, to the need to recognize patient-specific tumor antigens presented in the context of highly polymorphic major histocompatibility complex (MHC) or human leukocyte antigen (HLA) molecules.

Despite recent advances in the field of computational biology, *in silico* prediction of antigen-specific TCR sequences is still ineffective (3). Current TCR identification platforms rely on *in vitro* selection of antigen-specific T cells for subsequent TCR sequencing. These techniques often depend on the ability of individual T cells to bind peptide-MHC (pMHC) multimers or their capacity to proliferate and/or express activation markers after *in vitro* peptide stimulation (4). As such, these methods may introduce biases towards selection of high-affinity TCR sequences, which undergo more robust pMHC binding and proliferation. In addition, recent work has shown that T cells bearing antigen receptors of low and high affinity for a given antigen perform different functions. Indeed, while T cells bearing high affinity antigen receptors may, acutely, be more effective in their anti-tumor activity, they may also be more susceptible to inhibitory receptor mediated dysfunction and, potentially, off-target cross-reactivity (2, 5–12). Thus, it may be important to consider engineering T cells with a breadth of TCR affinities for optimal therapeutic efficacy.

In this context, antigen-specific T cell identification based on calcium (Ca^2+^) oscillations downstream of TCR signaling is an alternative approach with significant potential. TCR-dependent stimulation of T cells induces changes in intracellular Ca^2+^ concentration with temporal dynamics that contain information about TCR affinity (13–17). Furthermore, Ca^2+^ is a proximal readout of TCR activation, occurring within seconds of antigen receptor stimulation, limiting the potential selection biases induced by prolonged interaction with an antigen and expansion of a potentially limited number of clones. Genetic reporters for Ca^2+^ signaling (e.g., NFAT-GFP) have previously been used for the isolation antigen-specific TCR transduced T cells using a microfluidics system (18, 19), but their use for the discovery of antigen-specific TCR sequences from polyclonal T cells has not yet been achieved. The complexity of TCR-dependent Ca^2+^ signals and the possibility that TCR-independent processes impact intracellular Ca^2+^ levels are hurdles for its widespread use as a marker for TCR activation.

The use of supervised Machine Learning (ML) tools to process highly complex phenomena is revolutionizing approaches to clinical and fundamental research (20–23). Several studies have shown that these methods can be used to characterize T cell antigen-specificity from microscopy-based image datasets, by monitoring the interaction of T cells with antigen presenting cells (APC) or the autofluorescence changes that correlate with metabolic state (24–26). We propose to use these ML algorithms, trained to identify TCR-dependent Ca^2+^ fluctuations, to provide a prediction of antigen-specificity at the single cell level.

Here we present a proof-of-concept study for predicting T cell antigen-specificity based on intracellular Ca^2+^ dynamics. We took advantage of TCR transgenic T cells of known specificity, intracellular Ca^2+^ concentration indicator dyes, and simple imaging techniques to train and validate a ML model to accurately and efficiently predict antigen-specific T cells based on intracellular Ca^2+^ dynamics, which was then applied to polyclonal T cell responses. We show that convolutional neural networks (CNN) allow for efficient and accurate prediction of T cell activation from intracellular Ca^2+^ fluctuations at early time points, matching or surpassing other ML approaches. This method also demonstrates, for the first time, the feasibility of training algorithms on monoclonal TCR transgenic T cells, stimulated with model peptides, for the prediction of antigen-specificity in polyclonal T cell responses.

## Results

### In vitro T cell activation model to track intracellular Ca^2+^ dynamics

For the purpose of training an ML algorithm that predicts T cell antigen-specificity based on Ca^2+^ dynamics, we developed a simple imaging and analysis pipeline (Fig. 1A). To generate a widely-applicable and more physiologically-relevant *in vitro* system, we chose to develop an assay which uses peptide rather than pan-T cell stimulation (e.g., anti-CD3ε/CD28 antibodies, phytohaemagglutinin, etc.). With this method of stimulation, polyclonal T cells are poorly suited for training ML algorithms due to the very low frequency of antigen-specific T cells and the inability to know *a priori* the antigen-reactivity of individual cells. Indeed, standard ML training requires labeled ground-truth data as well as balanced datasets, with a similar number of positive and negative cells. We therefore used murine monoclonal OT-I and P14 TCR transgenic naïve CD8^+^ T cells in combination with Lipopolysaccharide (LPS)-matured bone marrow-derived dendritic cells (BMDC), loaded with chicken ovalbumin 257-264 (OVA) peptide (Supp. Fig. 1). CD8^+^ T cells are labeled with a ratiometric Ca^2+^ indicator dye, Indo-1, where the ratio between Ca^2+^-free and Ca^2+^-bound emission wavelengths is indicative of relative intracellular Ca^2+^ concentration. In all experiments, both OT-I and P14 T cells are co-cultured in the same well at a 1:1 ratio; a vital cytoplasmic stain, cell trace far red (CTFR), was used to label either OT-I or P14 T cells prior to Indo-1 staining in order to differentiate the two populations. Images for both Indo-1 and CTFR were captured over a period of two hours, beginning a few minutes after the start of the co-culture, to monitor intracellular Ca^2+^ dynamics. An *in silico* analysis pipeline was generated to automatically identify each cell, track it over time, measure the fluorescence of Indo-1 at each time point, and assign a genotype based on CTFR fluorescence. Thus, we can measure the dynamics of intracellular Ca^2+^ concentration for individual cells and know their antigen-specificity.

**Figure 1.**
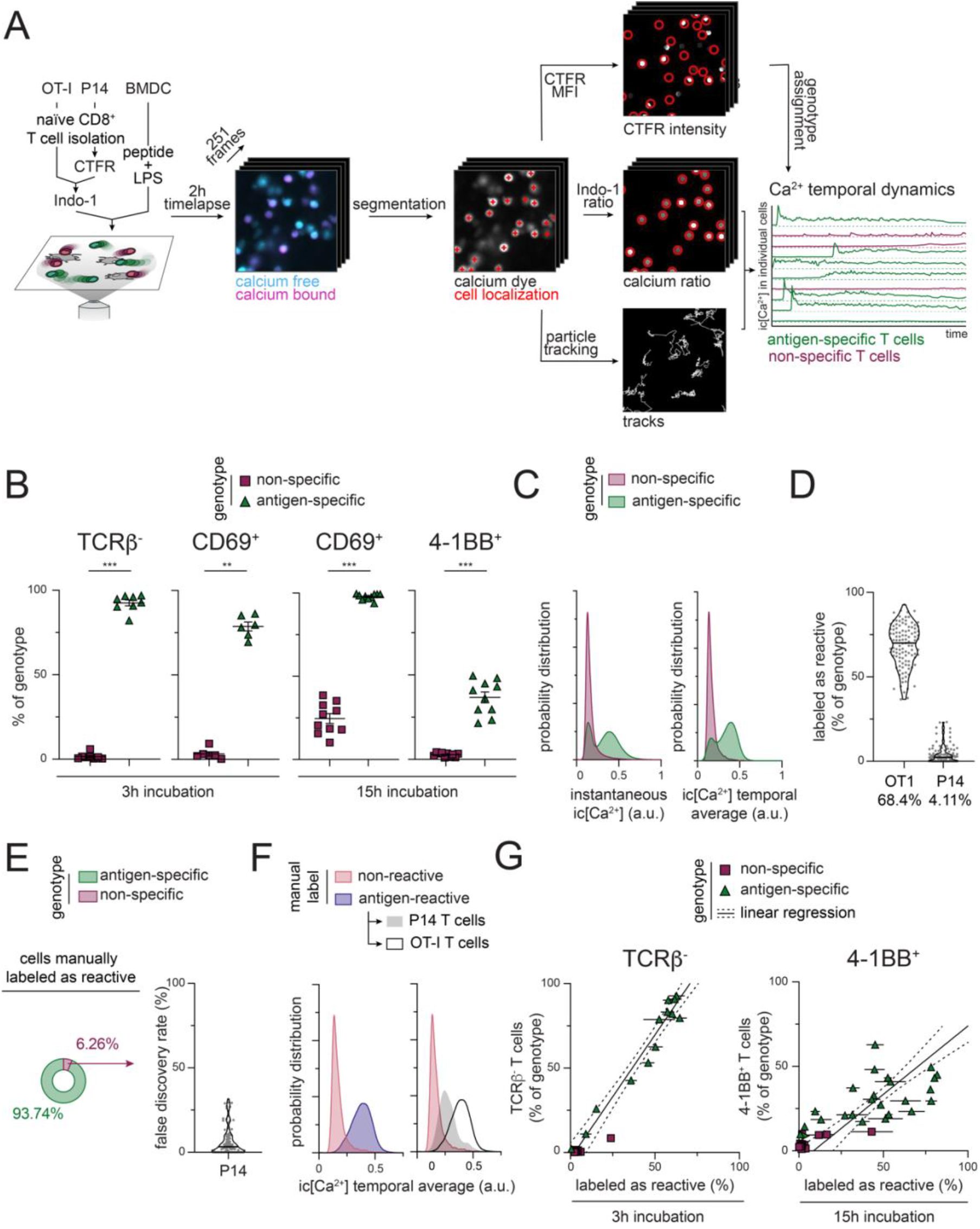
*In vitro* T cell activation model for the study of intracellular Ca^2+^ dynamics. (A) Schematic representation of the analysis pipeline. (B) Flow cytometry assessment of surface TCRβ down-regulation, CD69 expression, and 4-1BB expression, 3h or 15h after co-culture initiation. Error bars indicate standard deviation (SD). (n=6-10 independent culture wells respectively; ***=p<0.005; **=p<0.01 – Mann-Whitney U-test) (C) Probability distribution function of intracellular Ca^2+^ concentration for all timepoints (left) or the average Ca^2+^ concentration over the entire timelapse (right) according to antigen-specificity assignment. (n=7173 antigen-specific; n=7564 non-specific T cells) (D-G) The Ca^2+^ fluctuation of each cell was manually identified as antigen-reactive or non-reactive. (D) Frequency of T cells, expressed as a percentage of genotype, manually labeled as antigen-reactive. Horizontal lines in the violin plot show the median and numbers below show the average of the distribution. Individual fields of view are represented in gray. (n=111 fields of view) (E) Proportion of antigen-specific and non-specific cells among those manually labeled as antigen-reactive. A fraction of non-specific T cells have been manually labeled as antigen-reactive, making a false discovery rate for the manual labeling process of 6.26%. (F) Probability distribution function of the average intracellular Ca^2+^ concentration according to manual assignment and genotype. (n=5038 OT1 cells; n=347 P14 cells labeled as antigen-reactive; n=9351 cells labeled as non-reactive) (G) Frequency of cells manually labeled as antigen-reactive as a function of the frequency of cells downregulating surface TCRβ expression or expressing 4-1BB, as measured by flow cytometry, after 3h or 15h incubation for matching wells. For each independent culture well, data for both antigen-specific and non-specific T cells is shown, and the frequency of manual labeling for all 3 fields of view per well is averaged. Error bars indicate standard error of the mean (SEM) of manual labeling, full line shows linear regression of antigen-specific T cells, dotted lines show 95% confidence interval of regression. (TCRβ: n=13 independent wells, R^2^=0.9534, ρ=0.9764; 4-1BB: n=20 independent wells, R^2^=0.3767, ρ=0.6137)

For initial validation of the *in vitro* assay, we assessed T cell activation using well-established flow cytometric analysis of early (TCRβ downregulation and CD69 upregulation) and late (4-1BB expression) markers of T cell activation 3 and 15 hours after co-culture initiation. For these TCR transgenic models, we find TCRβ downregulation after 3 hours to be a high-fidelity read-out of activation (Fig. 1B and Supp. Fig. 2). CD69 and 4-1BB expression, however, show lower efficiency; they not only require more time to be robustly expressed, but there is also evidence of TCR-independent expression possibly driven by cytokines in the culture, as has been previously described (27–30). The distribution of Ca^2+^ concentration values, considering either all individual timepoints for all cells (left) or the average of each cell over the entire movie (right), shows a significant elevation in Ca^2+^ concentration only for antigen-specific T cells (Fig. 1C), while non-specific T cells display baseline intracellular Ca^2+^ concentrations. Altogether, these results show the appropriateness of this *in vitro* culture and analysis pipeline.

Not all antigen-specific T cells upregulate intracellular Ca^2+^ during the 2-hour imaging window. Because the development of an effective ML classifier, in principle, requires a high quality training dataset, these non-activated antigen-specific T cells could potentially interfere with the performance of a predictive model. Therefore, we manually labeled the Ca^2+^ signals of each cell in the dataset as antigen-reactive or non-reactive based on visual inspection of the Indo-1 fluorescence ratio. Four independent evaluators blindly classified each cell based on its relative Ca^2+^ levels over time, and a majority vote determined the final assignment of reactivity status; 68.4% of all antigen-specific T cells and 4.11% of non-specific T cells were labeled as antigen-reactive in the training datasets (Fig. 1D). This suggests that some Ca^2+^ fluctuation occurs in non-specific T cells, although this would not ultimately result in productive activation (Fig. 1B). Using manual labeling as a method to classify T cell antigen-specificity, we show a false discovery rate (FDR), i.e., the fraction of non-specific T cells within all cells labeled as antigen-reactive, of 6.26% (Fig. 1E). Two unimodal distributions of average Ca^2+^ concentration are observed based on manual assignment of cell status as antigen-reactive or non-reactive (Fig. 1F). However, non-specific cells manually labeled as antigen-reactive display an intermediate Ca^2+^ concentration distribution suggesting that, while difficult for human evaluators to differentiate from antigen-specific T cells, non-specific cells with intracellular Ca^2+^ levels above baseline have distinct Ca^2+^ fluctuation dynamics than that of *bona fide* antigen-specific T cells. Lastly, we show a positive correlation between the activation efficiency of each independent culture well, determined by manual assignment of Ca^2+^ traces and molecular activation markers measured by flow cytometry, both at early (TCRβ - Pearson correlation coefficient: ρ=0.976) and later (4-1BB - Pearson correlation coefficient: ρ=0.6137) timepoints (Fig. 1G), further validating the manual labeling process.

Increases in intracellular Ca^2+^ downstream of TCR engagement induce migration arrest (31). Given the importance of migration patterns in other approaches to identify T cell activation (24, 25, 32), we computed the speed of each cell averaged over the entire movie. We show that cells manually labeled as antigen-reactive are slower, on average, than those labeled as non-specific (Supp. Fig. 3A). Additionally, at timepoints where Ca^2+^ concentration is low on antigen-reactive T cells (prior to activation), the average and instantaneous velocity is identical to or above that of non-specific T cells (Supp. Fig. 3B & C).

### Deep-learning approaches perform better than conventional methods for the classification of T cell activation based on Ca^2+^ fluctuations

We systematically tested a non-exhaustive list of ML models which have been extensively used for the classification of 1-dimensional datasets. We divided the experiments into training and evaluation datasets, balancing the number antigen-specific and non-specific T cells, as well as the number of cells manually labeled as antigen-reactive and non-reactive. Despite having confirmed that CTFR staining of either OT-I or P14 did not affect the critical parameters of this co-culture setup (Supp. Fig. 4), we also balanced the amount of movies with both CTFR staining conditions to prevent models from learning specific features of either condition (Supp. Table 1). While the training datasets only contain co-cultures of TCR transgenic cells together with BMDC presenting OVA, the test datasets consist of co-cultures with either OVA or lymphocytic choriomeningitis virus (LCMV) gp33 (gp33-41) peptides to prevent overfitting and optimize the applicability of this model to a broader peptide repertoire. As expected, Ca^2+^ fluctuation of P14 T cells in gp33 co-cultures is upregulated as compared to their non-specific OT-I TCR transgenic counterparts (Supp. Fig. 5). Additionally, the manual labeling of the Ca^2+^ dynamics associated with gp33-stimulated T cell co-cultures shows a similar FDR to the OVA co-cultures (Supp. Fig. 5).

To benchmark the algorithms and choose an optimal architecture, we computed the efficiency (fraction of cells correctly predicted as antigen-specific) and FDR (fraction of mispredicted cells) for each model. Assuming that a subset of antigen-specific T cells will not be activated, the model efficiency is calculated by comparing the prediction to the manual label, rather than the genotype of the cell (OT-I or P14). On the other hand, to generate a model that best predicts whether a T cell is antigen-specific, the FDR compares the prediction to the genotype (i.e., non-specific T cells predicted as antigen-specific) for a measurement of accuracy. To compare models, the efficiency and accuracy metrics for both OVA and gp33 datasets are used to compute an *ad hoc* weighted performance metric, allowing a choice of the relative importance of accuracy over efficiency (see Methods); the model which maximizes this metric is chosen as optimal.

We first evaluated the use of a simple thresholding approach to classify each cell as either antigen-specific or non-specific based on relative intracellular Ca^2+^ concentration. Setting a first threshold dividing [Ca^2+^]^hi^ and [Ca^2+^]^low^ states, we computed the time each cell spent in each Ca^2+^ state. A second threshold is then set; any cell spending more than this amount of time in the [Ca^2+^]^hi^ state was classified as antigen-specific (Supp. Fig. 6A). For all possible pairs of thresholds, we computed the efficiency and accuracy of this method on the training dataset. The pair of thresholds which maximized the performance metric was then used to perform classification for the evaluation dataset (Supp. Fig. 6B). This approach has high efficiency (95.2%) but relatively poor accuracy (FDR=12.7%) for OVA co-cultures (Supp. Fig. 6C). Furthermore, this method is poorly applicable to co-cultures where gp33 is used; despite good accuracy (FDR=1.48%), a high percentage of antigen-specific cells were not predicted (70.4% efficiency), likely due to differences in average intracellular Ca^2+^ concentration between OVA- and gp33-specific TCR transgenic T cells (Supp. Fig. 6C & D).

To find a more suitable approach for identifying antigen-specific cells based on intracellular Ca^2+^ levels, we tested a multitude of models for accuracy and efficiency, going from simpler to more complex architectures and assessing the need of pre-processing and data-augmentation (Fig. 2A & Supp. Table 2). Using the performance metric to choose an optimal model, we show that deep learning algorithms were generally superior to other ML approaches. In particular, CNN-based architectures performed much better than any other method with this dataset (Supp. Fig. 6E), especially when the structure and training parameters are optimized (see Methods). The optimized CNN model using manually labeled cells as ground truth performs the best and is used for the rest of this study; it is referred to as optCNN_man_.

**Figure 2.**
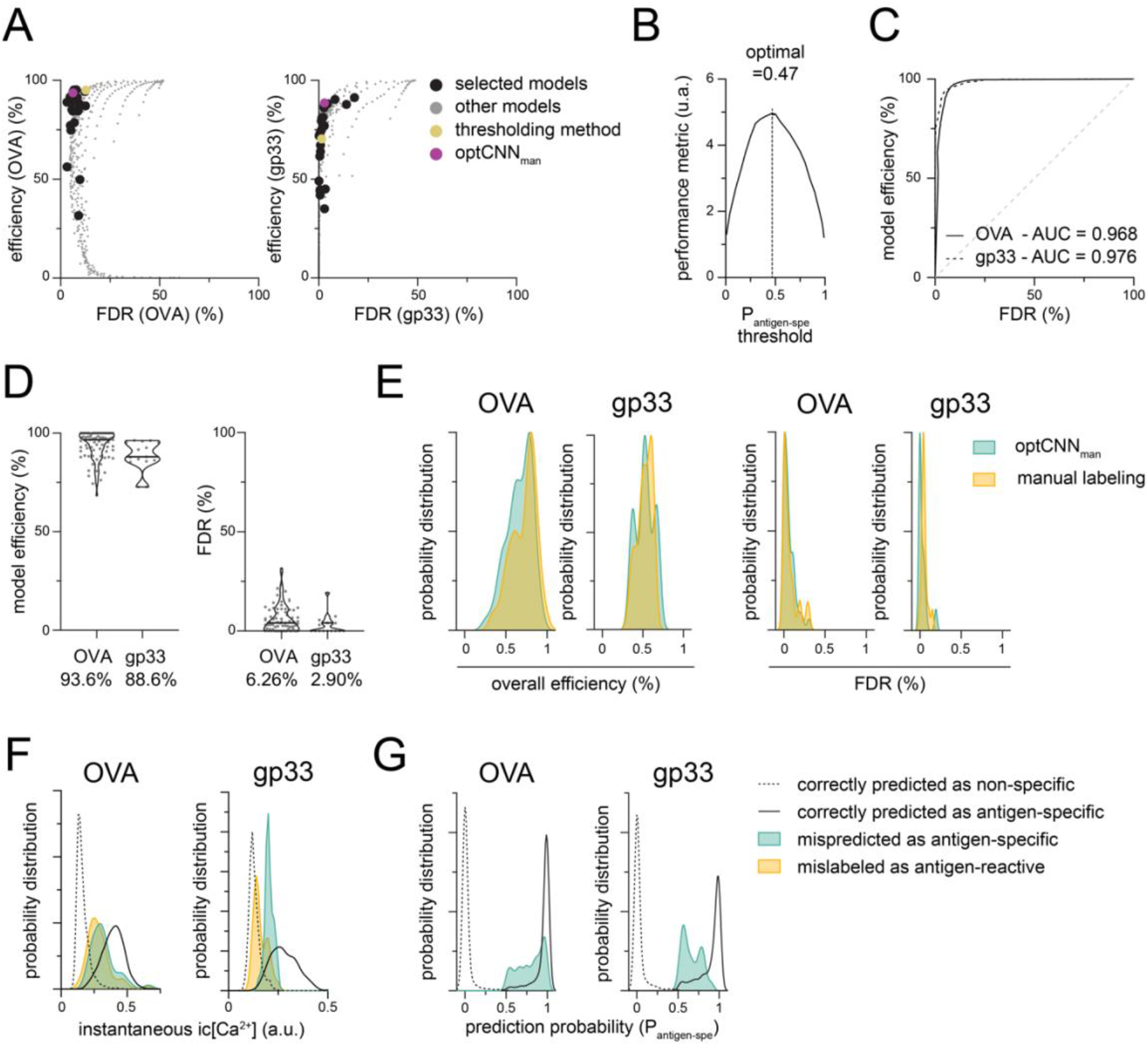
Convolutional neural networks allow for efficient and accurate classification of T cell activation based on intracellular Ca^2+^ dynamics. We conducted a systematic search through various machine-learning model architectures, evaluating their ability to predict antigen-specificity from Ca^2+^ fluctuations in individual cells. (A) Model efficiency (frequency of cells manually labeled as antigen-reactive predicted as antigen-specific) and false discovery rate (FDR - frequency of non-reactive T cells predicted as antigen-specific) of the prediction of all the machine learning algorithms tested, for both OVA (left) and gp33 (right) timelapses. Performance of the models detailed in Supp. Table 1 (empty circles and magenta) are plotted along with the performance of the thresholding approach (Supp. Fig. 6B-D). Gray dots represent other models generated during the systematic evaluation of ML structures with suboptimal performance, and are not detailed further. The optimal optCNN_man_ model, maximizing the performance metric, is shown in magenta. (B) Performance of optCNN_man_ monitored using the weighted performance metrics (see Methods) varying the threshold (t - see Supp. Fig. 6) on the probability of being antigen-specific (P_antigen-spe_) associated with each cell; cells above this threshold are classified as antigen-specific. (C) Receiver operating characteristic (ROC) curve of optCNN_man_ for both OVA (full line) and gp33 (dotted line) timelapses. Area under the curve (AUC) for both curves represents the performance of optCNN_man_ across all possible (P_antigen-spe_) thresholds. (D) Detailed performance of optCNN_man_ using the 0.47 prediction probability threshold. Horizontal lines in the violin plot show the median and numbers below show the average of the distribution. Individual fields of view are represented in gray. (n=73 OVA fields of view; n=15 gp33 fields of view) (E) Distribution of the overall efficiency (frequency of antigen-specific T cells predicted as antigen-specific) and FDR across all fields of view in the evaluation dataset, for optCNN_man_ and the manual labeling process. (n=73 OVA fields of view; n=15 gp33 fields of view) (F and G) Distribution of intracellular Ca^2+^ concentration (F) and prediction probability P_antigen-spe_ (G) of the non-specific cells mispredicted as antigen-specific. (n=174 cells in OVA co-cultures; n=32 cells in gp33 co-cultures)

This systematic approach revealed several important insights. Normalization of calcium concentration across independent experimental days (see Methods) is a critical factor, improving the efficiency of prediction of gp33 timelapses by over 28% (Supp. Table 2). Re-evaluating the thresholding method with data normalization shows an improved performance to non-normalized data but still lags behind CNN (Supp. Fig. 6F). Second, in this particular *in vitro* setup as opposed to other similar studies, we noticed that the use of positional data - using here the instantaneous speed of each cell - in conjunction with Ca^2+^ dynamics did not significantly improve classification (Supp. Table 2). Lastly, we observed that models trained with either the manual labels (T cells labeled as antigen-reactive vs labeled as non-reactive), the genotype (antigen-specific vs non-specific T cells) or a combination of both (antigen specific T cells labeled as antigen-reactive vs the rest) as ground truth, all perform relatively well (Supp. Table 2). When the training parameters are optimized (Supp. Table 3), all three models show a very similar performance (Supp. Fig. 6F), and their predictions overlap for 94.5% of the cells in the evaluation dataset (12,421 out of 13,145 cells) (Supp. Fig. 6G). Hence, it appears, for this application, that this architecture is not very sensitive to contamination of the dataset by negative (non-activated OT1) cells.

For each individual cell, optCNN_man_ provides a prediction probability; a threshold on this probability was used to determine the classification (Fig. 2B). The distribution of the probability of being antigen-specific (P_antigen-spe_) for all individual cells is bimodal, but classification of the rare cells that lie in between can drastically change the performance metrics. By varying the P_antigen-spe_ threshold, above which cells are predicted as antigen-specific, we show that the optimized architecture performs best when using the 0.47 P_antigen-spe_ threshold (Fig. 2B).

The receiver operating characteristic (ROC) curve shows the high sensitivity and specificity of the model with an area under the curve (AUC) of more than 0.95 (Fig. 2C). More specifically, optCNN_man_ has high efficiency for both OVA and gp33 co-cultures (94.1 and 88.6%, respectively) and low error rates (6.26 and 2.90%, respectively) (Fig. 2D). Furthermore, these predictions are similar to the predictions made by the human evaluators (Fig. 2E). Importantly, the intracellular Ca^2+^ concentration of the non-specific cells mispredicted as antigen-specific significantly overlaps with those of non-specific T cells manually labeled as antigen-reactive (Fig. 2F). Given the low prediction probability assigned to these cells (Fig. 2G), using a more restrictive threshold on P_antigen-spe_ would be expected to remove a significant number of the false positive predictions, at the cost of reduced efficiency.

### Biological validation of the Ca^2+^-based deep-learning algorithm to predict antigen-specificity

We next sought to validate optCNN_man_ using an alternative biological approach. By altering the parameters of the BMDC:T cell co-culture (e.g., number of antigen presenting BMDC, total number of BMDC, etc.) to modulate activation efficiency, we investigated how efficiently optCNN_man_ can predict activation in sub-optimal conditions. In theory, the lower absolute number of peptide-loaded APC would lead to an increase in the time required for T cells to find cognate antigen, which translates, given the short imaging window, into a smaller fraction of activated cells. Using TCRβ downregulation and 4-1BB expression after 3h and 15h of incubation, respectively, we show a positive correlation between the cells predicted as antigen-specific with these markers (Fig. 3A). Furthermore, manipulating the ratio of antigen-presenting to non-presenting BMDC in individual co-culture wells, we show, as anticipated, a dose-dependent relationship between the frequency of T cells predicted as antigen-specific and antigen availability (Fig. 3B). Similarly, the overall BMDC number in the co-culture also leads to a reduction in the number of T cells predicted as antigen-specific (Fig. 3B).

**Figure 3.**
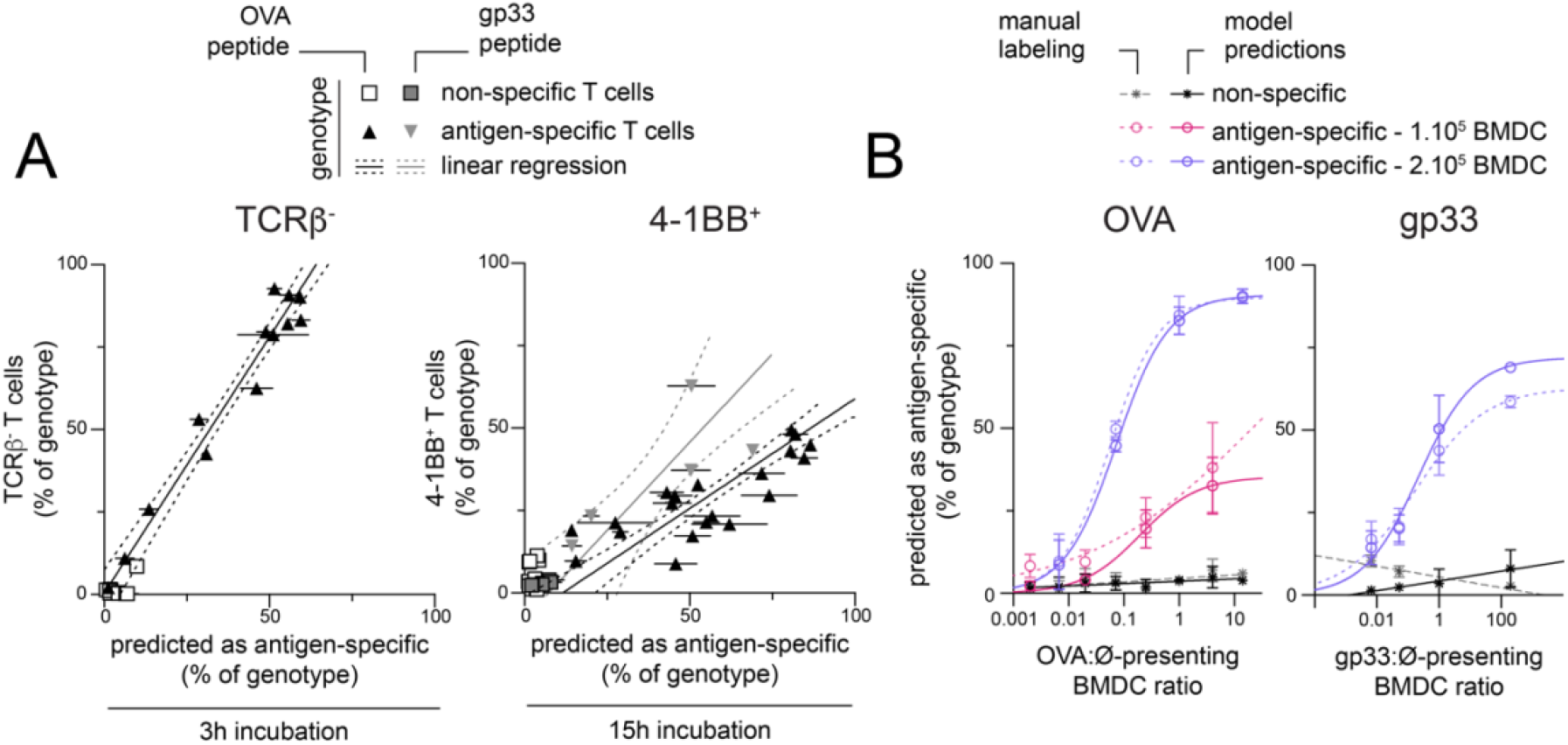
Biological validation of antigen-specificity predictions based on intracellular Ca^2+^ dynamics. Critical parameters of the co-culture (e.g., number of antigen presenting BMDC, total number of BMDC, etc.) are modulated before co-culture; optCNN_man_ is then used to predict the frequency of antigen-specific T cells. For each independent culture well, the prediction percentage is averaged over three fields of view; wells are then harvested and analyzed by flow cytometry for expression of selected activation markers. (A) Correlation between ML prediction and the frequency of cells downregulating surface TCRβ expression or expressing 4-1BB, as measured by flow cytometry, after 3h and 15h incubation, respectively. Error bars indicate SEM. Linear regression of the antigen-specific conditions is shown (full line) with 95% confidence error (dotted lines). (TCRβ: n=12 independent wells, R^2^=0.959, ρ=0.989; 4-1BB & OVA peptide: n=20 independent wells, R^2^=0.651, ρ=0.807; 4-1BB & gp33 peptide: n=5, R^2^=0.581, ρ=0.762) (B) Percentage of antigen-specific and non-specific T cells predicted as antigen-specific as a function of the ratio of antigen-presenting to non-presenting BMDC. Percentage of cells predicted as antigen-specific is averaged for the 3 fields of view of each co-culture. Full lines and dotted lines represent a sigmoidal curve fitted to the data. The non-specific group pools data from both 1×10^5^ and 2×10^5^ BMDC conditions. Error bars indicate SD.

Because it is not possible to know *a priori* the antigen-specificity of individual naïve polyclonal CD8^+^ T cells, the validation of the model on polyclonal responses to antigenic peptides is challenging without extensive experimental confirmation. Thus, we used a mixed lymphocyte reaction (MLR) which typically leads to a larger fraction of T cells being activated in a polyclonal fashion, as compared to antigen-specific T cells. We co-cultured C57BL/6J CD8^+^ T cells with MHC-matched C57BL/6J (autologous) or MHC-mismatched BALB/c (alloreactive) BMDC (Fig. 4A). Using molecular markers of activation measured by flow cytometry after 3h and 20h, we show that alloreactive culture conditions lead to a higher fraction of T cells expressing activation markers than in autologous conditions. Although visible as soon as 3h, monitoring of activation by flow cytometry is much more efficient after 20h of co-culture (Fig. 4B). While cell surface TCRβ downregulation was a very specific marker for monoclonal T cell activation, it is not an obvious marker of T cell activation in the MLR setting (Fig. 4B). Using optCNN_man_ to predict T cell activation based on Ca^2+^ fluctuations in this polyclonal system, we also show that T cells cultured in alloreactive culture conditions have a higher frequency of cells predicted as antigen-specific than when T cells are cultured with autologous BMDC (Fig. 4C). Additionally, there is a strong correlation between the ML predictions and the flow cytometry markers of activation, particularly after 20h of culture, confirming the accuracy of prediction (Fig. 4D). The distribution of intracellular Ca^2+^ concentrations in polyclonal T cells predicted as antigen-specific in the MLR is much wider than that of the monoclonal T cell populations used earlier (Fig. 4E); this may be due to the wider range of affinities in the polyclonal T cell repertoire and the difference in TCR:pMHC binding biomechanics (33).

**Figure 4.**
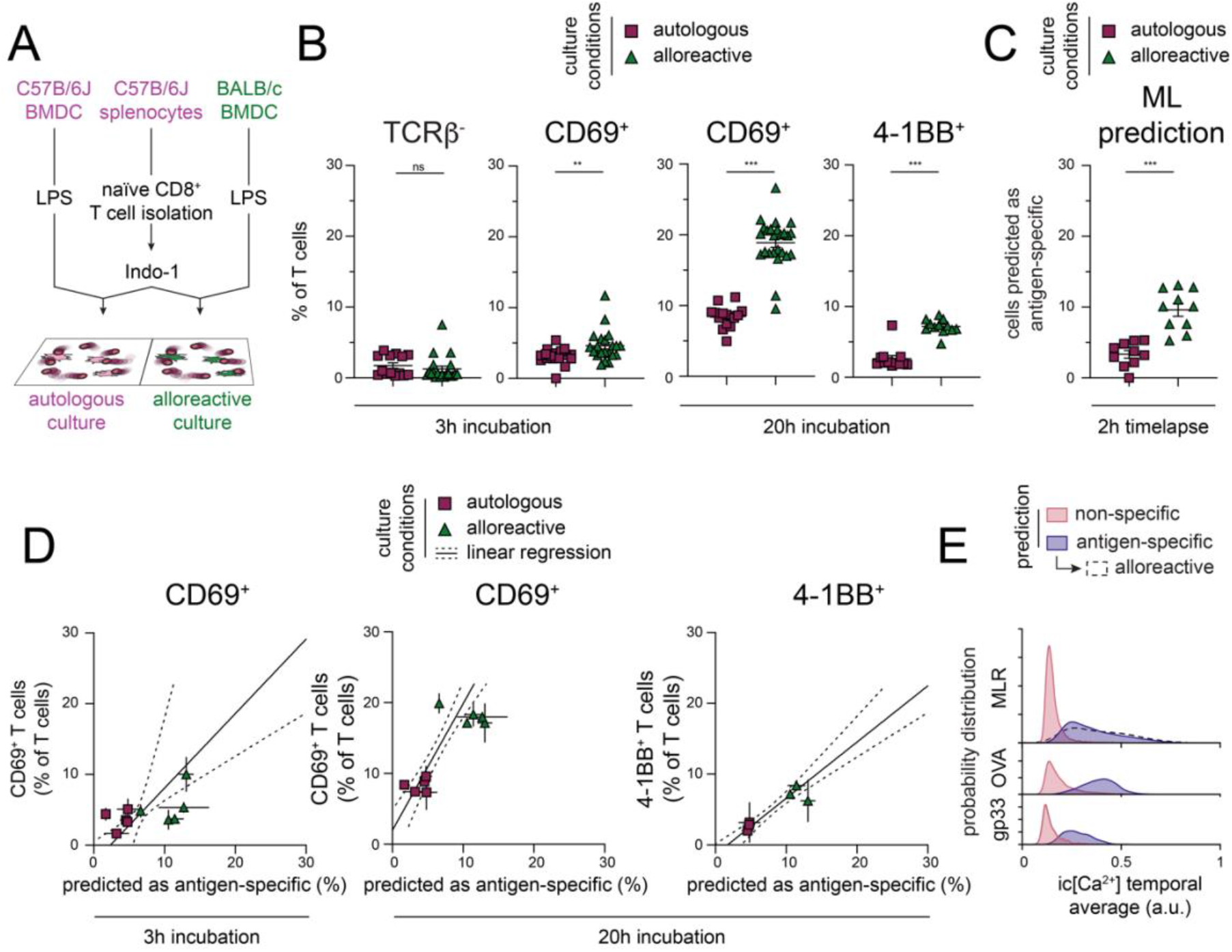
Deep-learning models predict polyclonal T cell responses. (A) Schematic representation of the mixed lymphocyte reaction (MLR) setup. Purified C57B/6J naïve CD8^+^ T cells, stained with Indo-1, were overlaid on MHC-matched C57B/6J BMDC (autologous conditions) or MHC-mismatched BALB/c BMDC (alloreactive condition). (B) Flow cytometry based measurement of TCRβ, CD69 and 4-1BB expression, 3h or 20h after co-culture as for alloreactive and autologous co-cultures. Error bars show SD. (n=4-10 culture wells; ***=p<0.005 – Mann-Whitney U-test) (C) Frequency of cells predicted as antigen-specific (D) Correlation between ML prediction of antigen-specificity and the frequency of cells expressing CD69 or 4-1BB (B), as measured by flow cytometry, after 3h and 20h incubation respectively. Each point is the average of the prediction of 3 independent fields of view and the average of 2-6 independent culture wells for flow cytometry data. Error bars indicate SEM of prediction and flow cytometry data. Linear regression is shown (full line) with 95% confidence error (dotted lines) and its correlation coefficient (R^2^) and slope. (n=3-5 independent experiments) (E) Probability distribution of the average Ca^2+^ concentration over the whole timelapse for all the cells according to the prediction by optCNN_man_ and culture condition, comparing prediction for MLR cultures and monoclonal T cell culture with OVA and gp33 antigens (replotted from Fig. 1 and Supp. Fig. 5). (MLR: n=3219 non-specific cells, n=541 antigen-specific cell; OVA: n=7266 non-specific cells, n=3333 antigen-specific cells; gp33: n=1904 non-specific cells, n= 642 antigen-specific cells)

Altogether, these data show the applicability of optCNN_man_, trained on monoclonal T cells responding to a single high-affinity peptide, for the prediction of responses to additional peptides as well as polyclonal T cell responses. Thus, simple models of T cell activation can be used to train ML architectures to recognize general features of Ca^2+^ fluctuation which are common to T cell responses across a wider range of TCR-pMHC affinities.

## Discussion

The rapid identification of antigen-specific T cells from naïve polyclonal T cells presents a unique challenge due to the lack of reliable early markers of TCR-specific T cell activation prior to proliferation. Here, we demonstrate the feasibility of using the time-dependent fluctuation of intracellular Ca^2+^ concentration in individual T cells, a TCR-proximal read-out, as a means to identify their antigen specificity. While increases in intracellular Ca^2+^ may not be strictly TCR-specific, we propose that ML algorithms, trained on T cells of known specificity and activated in an antigen-specific manner can learn the features of Ca^2+^ fluctuation associated with TCR-pMHC engagement. We show that, once trained on monoclonal T cell responses, this model can be applied to predict activation of polyclonal T cells.

As compared to other methods of antigen-specific T cell identification, we suggest that the monitoring of intracellular Ca^2+^ signaling is a much faster and simpler approach to the identification of antigen specific T cells. Although they do not require T cell stimulation, thereby bypassing the delay in the modulation of activation marker expression, multimer pMHC-based enrichment requires the engineering of a new reagent for each peptide and individual peptide/MHC combination, some of which may be problematic to manufacture (34). Early T cell activation is also challenging to measure by flow cytometry, due to the absence of appropriate markers. We show that, while early (3h) surface TCRβ downregulation and CD69 expression are specific markers of monoclonal T cell activation in the *in vitro* model shown here, they were not useful in the more physiologically-relevant polyclonal MLR cultures. 4-1BB, although more specific, appears to be upregulated much later and only maximally after some proliferation has occurred, limiting its usefulness for early isolation of antigen-specific T cells and limiting biased expansion of high-affinity T cell clones. In comparison, increases in intracellular Ca^2+^, a very early indicator of TCR signaling pathway, allows the quick - within a 2h timeframe - and accurate identification of antigen specific T cells. The simple nature of the co-culture setup, commonly used for antigen-specific T cell activation, allows for flexibility in the target antigen (e.g., cancer antigens) loaded and MHC/HLA-restriction, by varying the source of APCs.

Recent studies have also investigated the use of imaging-based technologies and ML for the identification of antigen specificity, investigating either the dynamics of interaction between antigen-specific T cells and APCs or the changes in metabolic state associated with pan-T cell activation. (24–26). With an AUC of more than 0.95, the performance of the approach presented in this paper is similar to, if not superior to these previously published studies. Additionally, the stimulation of naïve CD8^+^ T cells with peptide-loaded BMDC, rather than pan T cell stimulation, ensures that our ML models learn features of Ca^2+^ fluctuation which are generated during physiological TCR engagement.

In terms of experimental complexity, the use of simple, cheap and widely accessible fluorescent dyes and labware for conventional fluorescence microscopes are the only requirements and represent a low cost-of-entry for using this technology for downstream applications. Extraction of Ca^2+^ fluctuations from these movies and the training of the 1-dimensional ML network also has the major advantage of requiring very little computing power and can be replicated with any desktop computer. A few studies have demonstrated the use of microfluidics, micropipettes and/or microraft apparatus for the isolation of antigen specific T cells which may also be challenging to manufacture and are relatively low throughput technologies (18, 35–38).

Here we made use of naïve T cells for the evaluation of Ca^2+^ fluctuations. Naïve T cells, as opposed to antigen-experienced T cells, are not restricted in their TCR repertoire as may be the case after clonal expansion and may allow for the identification of TCR with a wider range of affinities for a peptide of interest. Furthermore, naïve and antigen-experienced T cells, and even different antigen-experienced T cell subsets, display distinct Ca^2+^ fluctuation patterns in response to TCR stimulation (39, 40). The use of purified naïve T cells, although themselves heterogeneous in nature (41), should allow for more homogenous and reproducible Ca^2+^ response to TCR stimulation, compared to antigen-experienced T cells.

In this proof-of-concept study, we demonstrated that performant ML algorithms can be trained on Ca^2+^ fluctuations in activated monoclonal T cells to predict polyclonal T cell responses, using a limited amount of data (∼10,000 cells). Substantially increasing the size of the dataset with additional timelapses or through AI-assisted methods (42–44) may further improve model performance, especially when it comes to differentiating the distinct pattern of Ca^2+^ fluctuation associated with non-specific T cells mispredicted as antigen-specific from *bona fide* antigen-specific T cells. Furthermore, given that many factors of TCR signaling, such as antigen avidity and costimulation, induce distinct patterns of intracellular Ca^2+^ concentration variation (13, 14, 45, 46), the use of Ca^2+^ fluctuations from monoclonal T cells, following TCR stimulation with peptide-MHC of varying affinity, should enable training of ML models to recognize specific features of Ca^2+^ fluctuation associated with low versus high affinity binding to antigen. It has already been shown that an ML model, trained on the dynamics of cytokine release by T cells following stimulation over several weeks, can predict antigen affinity of individual T cells (47). The use of intracellular Ca^2+^ dynamics would fast-track and simplify this approach.

Combining ML approaches for the identification of antigen-specific T cells with technologies that allow for the labeling of individual cells with high specificity based on observable characteristics will allow for the isolation of T cells of interest for downstream single cell TCR-sequencing (48); giving access to clinically relevant TCR sequences for use in adoptive therapy. Particular attention will need to be paid to the compounding of errors at the various steps of the pipeline (tracking, ML prediction & barcoding) to avoid contamination by non-specific TCR sequences, which would increase the time required for downstream biological validation of identified TCR sequence. However, the advent of fast and reliable *in vitro* and *in silico* pipelines for TCR-pMHC screening mitigates this risk (49, 50). Furthermore, the possibility to identify TCR sequences with a specific affinity, fine-tuned either for acute antitumoral activity (high affinity) or for longer lasting, broader immune-surveillance, with reduced side effects (lower affinity) could facilitate improvement in the quality of care to patients requiring adoptive T cell therapy (7, 9).

## Materials and Methods

### Mice

C57BL/6J and BALB/c mice were purchased from the Jackson Laboratory (Bar Harbor, ME, USA). C57BL/6J-Tg(OT-I)-Rag1<tm1Mom> (OT-I) mice were obtained through the National Institute of Allergy and Infectious Diseases Exchange Program, National Institutes of Health (Bethesda, MD, USA) (51, 52). P14 TCR Tg mice were provided by Dr. Martin Richer (McGill University, Montreal, Canada) and crossed onto a TCRα KO (The Jackson Laboratory, Stock# 002116) background (53, 54). All mice were bred and maintained in specific pathogen-free animal facilities at the Maisonneuve-Rosemont Hospital Research Centre. Both male and female mice 6– 12 weeks of age were used. All animal protocols have been approved by the Animal Care Committee at the Maisonneuve-Rosemont Hospital Research Centre. Experiments were performed in accordance with the Canadian Council on Animal Care guidelines.

### In vitro T cell co-culture assay

1×10^6^ bone marrow cells, harvested from the indicated mice, are plated in 6 well adherent plates for 7 days in 4mL of 10% FBS (HyClone, Cat#SH30396.03), 100mM HEPES (Multicell, Cat#330-050-EL), 100 IU Penicillin/Streptomycin (Multicell, Cat#450-201-EL), 1mM Sodium Pyruvate (Multicell, Cat#600-110-EL), 0.1 mM MEM non-essential amino acids (Gibco, Cat#11140-050), 2mM L-glutamine RPMI (Multicell, Cat#350-000-CL). Culture medium is supplemented with 1000U/well murine GM-CSF (Biolegend, Cat#576302) and a predetermined dose of P815-IL4 supernatant. 2mL of supplemented media is replaced after 2 and 3 days of culture. At day 6, 4μM of OVA (OVA 257-264, Anaspec Inc., Cat#AS-60193-5) or gp33 (gp33-41, Anaspec Inc., Cat#AS-61669) peptide and 1μg/mL LPS is added to the culture. At days 7, 8, or 9, BMDC are harvested from culture and enriched using a 14.7% Histodenz (Sigma, Cat#D2158) gradient.

For T cell isolation, cellular suspension is harvested via physical dissociation from OT-I and P14 spleen and lymph nodes. Naïve CD8^+^ T cells are further isolated using a magnetic enrichment kit according to manufacturer specifications (StemCell, Cat#19858). OT1 or P14 cells are stained with 2μM Cell Trace Far Red (Invitrogen, Cat#C34572) at 10^6^ cells/mL for 15 minutes at 37°C and rested 15 minutes at 37°C, 5% CO_2_ before being pooled. The cell suspension is then stained with 10μM Indo-1 for 30 minutes at 37°C and rested 30 minutes at 37°C, 5% CO_2_. The isolation and staining procedure is identical for C57B/6J and BALB/c naïve CD8^+^ T cells, but the T cells are kept separate at all times and are not stained with CTFR.

Right before imaging, unless otherwise specified, 2×10^5^ BMDC and 2×10^5^ T cells are pooled in phenol red-free imaging medium (10% Fetal Bbovinne Prodcut (HyClone, Cat#SH30109.03), 100 IU Penicillin/Streptomycin) and plated onto fibronectin-coated (2μg/cm^2^, Sigma, Cat#F2006) 18 well slides (IBIDI, Cat#81816) for imaging. Wide field epifluorescence images of Indo-1 (405nm and 447nm) and CTFR (698nM) are captured every 30 seconds (Indo-1) or 10 minutes (CTFR) for 2 hours on a Nikon Eclipse Ti2, under a stage top incubator, with a mercury lamp illumination. After imaging, slides are kept in an incubator before harvesting for flow cytometric analysis.

### Flow cytometric analysis

Flow cytometry analysis of mouse surface antigens was performed with the following Abs: Anti-CD3 (145-2C11, Cat#100328, 1:100), -CD8α (53-6.7, Cat#100714, 1:400), -CD11c (N418, Cat#117308, 1:1600), -CD45.2 (104, Cat#109822, 1:100), -CD80 (16-10A1, Cat#104722, 1:400), -I-A/I-E (M5/114.15.2, Cat#107630, 1:1600), -TCRβ (H57-597, Cat#109224, 1:100), -NK1.1 (PK136, Cat#108706, 1:200), -CD19 (6D5, Cat#115505, 1:200), -CD11b (M1/70, Cat#101206, 1:200), -TCRgd (N418, Cat#117306, 1:200), -CD44 (IM7, Cat#103012, 1:200), -CD69 (H1.2F3, Cat#104522, 1:100), -CD137 (17B5, Cat#106105, 1:200), and Zombie Aqua (Cat#423101) or Green fixable viability dye (Cat#423111) (BioLegend). Staining was performed for 20 min at 4 °C. Flow cytometry analyses were performed on a LSR Fortessa X20, and data were analyzed using FlowJo software (BDBiosciences).

### Prediction of T cell antigen-specificity

All the code used for the prediction of T cell specificity was coded using Matlab (MathWorks) and is available upon request.

### In silico analysis pipeline

From raw images, segmentation and tracking of T cells are made by adapting previously published methods (55–57). Briefly, in each frame, the centroid of each T cell is localized from the sum of the Indo-1 images using a combination of edge detection and watershed segmentation methods. Based on the position of all cells at each timepoint we calculate individual T cell trajectories using particle tracking-based methods, adapted for our particular application. Filtering of tracks based on length (at least half of the duration of the movie) ensures that all cells in a timelapse are independent from each other. The distance traveled by a cell between 2 frames is reported as instantaneous speed. Manual quality control on the tracking is made to remove mistracked cells.

For each cell, at each timepoint, the fluorescence intensity of both Indo-1 emission wavelengths is calculated by integrating pixel intensity values in a disk (60% of the cell’s diameter) around the centroid. The intensity background (calculated locally for each cell) is subtracted to the fluorescence intensity before calculating the ratio of both wavelengths (405nm/447nm). Assembling the ratio for each cell across all timepoints allows us to generate the intracellular Ca^2+^ dynamics. For genotype assignment, the fluorescence of CTFR is obtained 12 times throughout the imaging period for each cell; cell type is assigned if at least 8 timepoints the cell appears positive or negative for CTFR.

For manual labeling, four independent evaluators were shown the intracellular Ca^2+^ dynamics of all cells and were asked to classify them as antigen-reactive or non-reactive. A majority vote between evaluators was used to determine the final label of each cell.

### Training and evaluation of the machine learning algorithms

For all individual models, training was made on the training dataset using the 1D intracellular Ca^2+^ signal as input and the genotype of each cell, their manual label, or a combination of both (antigen-specific T cells manually labeled as antigen-reactive) as ground-truth. When indicated, Ca^2+^ dynamics were complemented with either the derivative of the Ca^2+^ dynamics, approximated by the absolute value of the difference in Ca^2+^ levels between two consecutive timepoints, or with the instantaneous cell speed, approximated by the euclidean distance between the same cell at two consecutive timepoints. Evaluation of the model was made on the evaluation dataset by predicting antigen-specificity for each individual timelapse (field of view).

### Performance metric

For each timelapse the model efficiency, overall efficiency and false discovery rate were measured as follow:

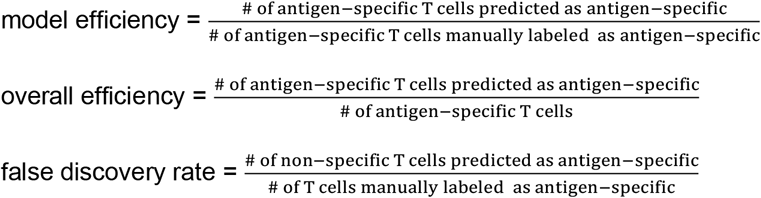

All throughout, the performance metric (pM) uses the average model efficiency (eff) and the average FDR across all timelapses in the evaluation dataset. The optimal model is the one that maximized the formula:

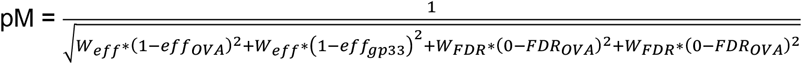

*w*_*eff &*_ *w*_*FDR*_ are variable parameters that modulate the importance attributed to the error rate vs the efficiency for each particular application. For this study, we use *w*_*eff*_=1 and *w*_*fdr*_=5.

### Data normalization

When indicated, Ca^2+^ concentration of each cell was normalized to the average value of ‘resting’ intracellular Ca^2+^ concentration. Briefly, for each timelapse, two Gaussian distributions are fitted to the probability distribution function of intracellular Ca^2+^ calcium concentration. The mean of each Gaussian distribution is used as the average value Ca^2+^ concentration in the low ([Ca^2+^]^low^) and high ([Ca^2+^]^hi^) state for this movie. For each cell at each timepoint, we divide its Ca^2+^ concentration by the average [Ca^2+^]^low^ value to compute the ‘fold-change’ of Ca^2+^ concentration over the ‘resting’ state, as a means to reduce inter-experiment variability.

### Data augmentation procedure

When indicated, data augmentation was performed by artificially generating *in silico* new Ca^2+^ fluctuations from real *in vitro* fluctuations. This is achieved by a combination of repeating existing data, adding noise to the existing data, shifting the start of Ca^2+^ fluctuation forward or backwards (in time) and increasing or decreasing the levels of the Ca^2+^ in the existing data.

### Hyperoptimization

For each parameter to be optimized, i.e., number of neurons, kernel size, optimizer, and mini batch size, a range of possibilities was determined according to commonly used values for that parameter in the literature. A model was trained and evaluated as previously described for each combination of these four parameters, across all the ranges. All the models were then evaluated using the weighted performance metric; the hyperoptimized model is the one maximizing the performance metric.

## Acknowledgments

We would like to thank the staff from the flow cytometry and animal facilities at the Maisonneuve-Rosemont Hospital Research Centre for their technical help. We are grateful to Johannes Textor and Shabaz Sultan from Radboud University, Netherlands as well as all members of the lab for all the fruitful discussions and their help with the manual labeling process. We would also like to thank Javier Mazzaferri, Joannie Roy, and Nicolas Desjardins Lecavalier for their previous work on the particle tracking algorithms.

This work was supported by the FRQNT (253380 to S.C. and H.J.M.) and NSERC Discovery grants (RGPIN-2021-03330 to S.C. and RGPIN-2019-05053 to H.J.M.). S.T. was supported by post-doctoral training scholarships from the Cole Foundation and the FRQS. H.J.M. is a senior scholar of the FRQS.

