## Supplementary Figures and Tables for "Predicting T cell activation based on intracellular calcium fluctuations"

### **Supporting Information for**
Predicting T cell activation based on intracellular calcium fluctuations

Sébastien This [1,2], Santiago Costantino [1,3,*], Heather J. Melichar [1,4,*]

*Santiago Costantino

*Heather Melichar

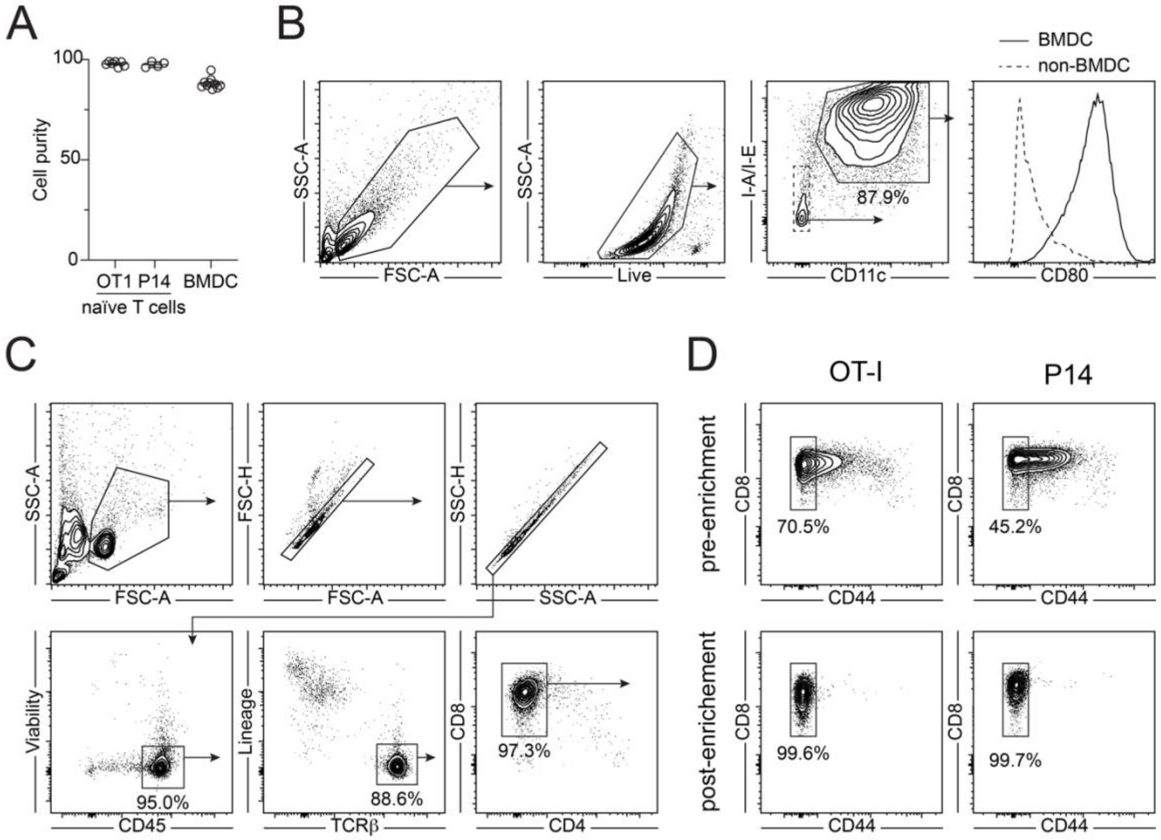


**Fig. S1.** Cell purities for the *in vitro* co-culture system (A) Purity of CD8$^+$ naïve T cells (Live Lin$^-$ CD45$^+$ TCRβ$^+$ CD8$^+$ CD44$^-$) and BMDC (Live CD11c$^+$ I-A/I-E$^+$) used in the co-culture assay as measured by flow cytometry. Error bars show SD. (n=7-11; Lin$^-$ = CD19$^-$ CD11c$^-$ NK1.1$^-$ CD11b$^-$ TCRγδ$^-$) (B) Gating strategy for the measure of BMDC purity and representative plot of CD80 expression on BMDC and non-BMDC as control. (C-D) Gating strategy to evaluate the purity of naïve CD8$^+$ T cells (Live Lin$^-$ CD45$^+$ TCRβ$^+$ CD8$^+$ CD44$^-$) for OT-I and P14 cells as measured by flow cytometry. (D) Representative plots of naïve CD8$^+$ T cells frequency before and after magnetic enrichment of naïve OT-I and P14 T cells.

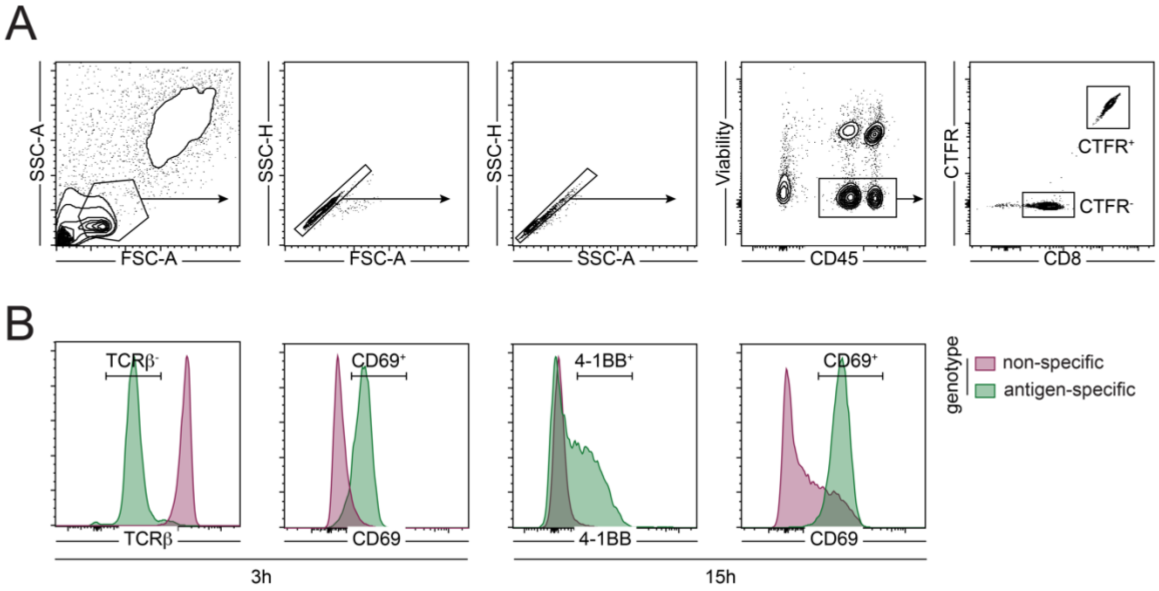


**Fig. S2.** Evaluation of monoclonal T cell activation by flow cytometry. (A) Gating strategy used for the identification of CTFR$^+$ and CTFR$^-$ CD8 T cells following co-culture. The genotype is determined based on which cells (OT-I or P14) were stained with CTFR prior to co-culture. (B) Representative distribution of TCRβ, CD69 and 4-1BB expression in antigen-specific and non-specific T cells, after 3 and 15 hours of co-culture as indicated.

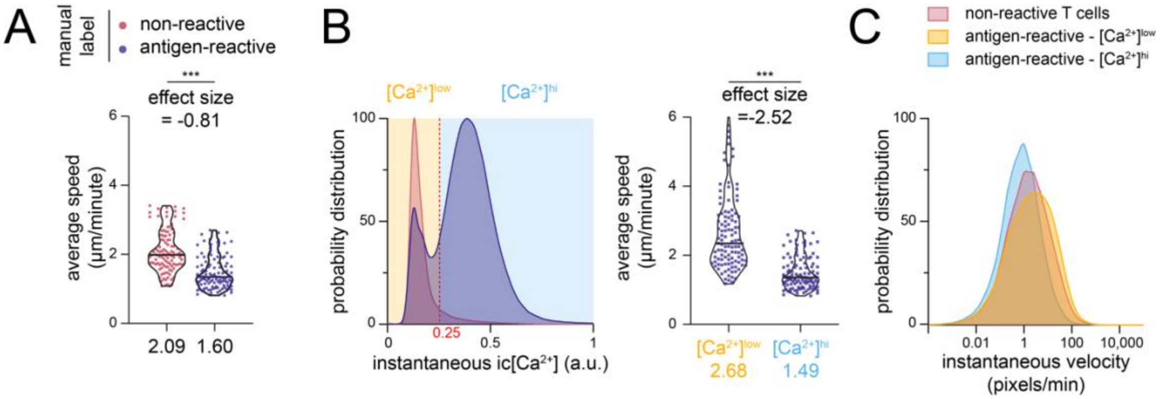


**Fig. S3.** Calcium signaling is associated with reduced T cell speed in the *in vitro* co-culture assay. (A) Quantification of average speed across the entire timelapse for all T cells based on manual labeling assignment. The speed of all cells with the indicated label is averaged in each field of view and is plotted here. Horizontal lines in the violin plot show the median and numbers below show the average of the distribution. Effect size between both distributions is computed using the left condition as control. (B) For each timelapse, an Otsu threshold (red line; threshold=0.25) on the instantaneous $Ca^{2+}$ concentration of all cells is used to separate $[Ca^{2+}]^{low}$ from $[Ca^{2+}]^{hi}$ states. The average speed for all cells at timepoints where they are in either $Ca^{2+}$ states is plotted on the right. (C) Distribution of instantaneous velocity in all cells for all timelapses according to the antigen-specificity and $Ca^{2+}$ concentration status. (n=111 fields of view; n=7173 antigen-reactive; n=7564 non-reactive T cells; ***=p<0.005 – Mann-Whitney U-test)

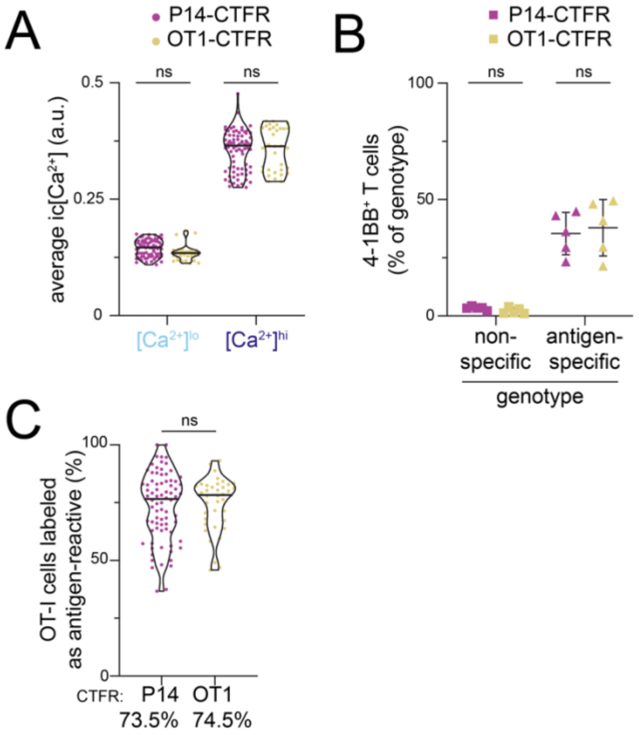


**Fig. S4.** CTFR on OT1 or P14 does not affect activation. (A) For each timelapse, two Gaussian distributions were fitted to the distribution of $Ca^{2+}$ concentration across all cells (Fig. 1C). The mean of these distributions is used as a surrogate for the mean intracellular $Ca^{2+}$ concentration of the $[Ca^{2+}]^{hi}$ and $[Ca^{2+}]^{low}$ populations. (B) Flow cytometric measurement of 4-1BB expression after 15h co-culture. Error bars show SD. (n=5 culture wells) (C) Frequency of OT-I, expressed as frequency of genotype, manually identified as antigen-reactive by four independent evaluators. Horizontal lines in the violin plots (A and C) show the median and numbers below show the average of the distribution. (n=78 P14-CTFR fields of view; n=38 OT1-CTFR fields of view; ns=p>0.05 – Mann-Whitney U-test)

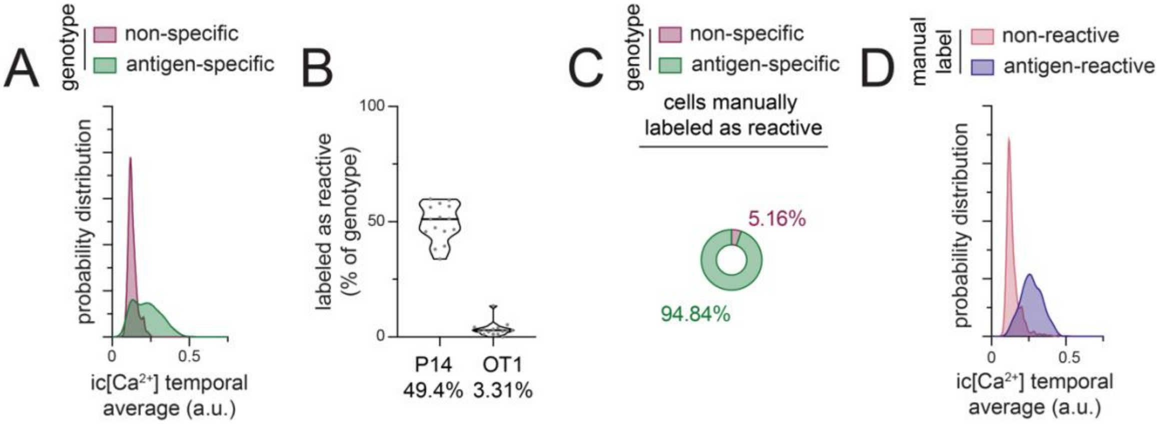


**Fig. S5.** Validation of the specificity of the co-culture model when using the gp33 peptide. (A) Probability distribution of the average $Ca^{2+}$ concentration over the whole timelapse according to antigen-specificity assignment. (n=1108 antigen-specific cells; n=1168 non-specific cells). (B) Frequency of T cells, expressed as percentage of genotype, manually labeled as antigen-reactive. Horizontal lines in the violin plot show the median and numbers below show the average of the distribution. Individual fields of view are represented in gray. (n=15 fields of view). (C) Detailed composition of cells manually labeled as antigen-reactive. Some of these are non-specific, making a false discovery rate for the manual labeling process of 5.16%. (E) Probability distribution of the average intracellular $Ca^{2+}$ concentration according to manual assignment (n=594 cells labeled as antigen-reactive; n=1682 cells labeled as non-reactive).

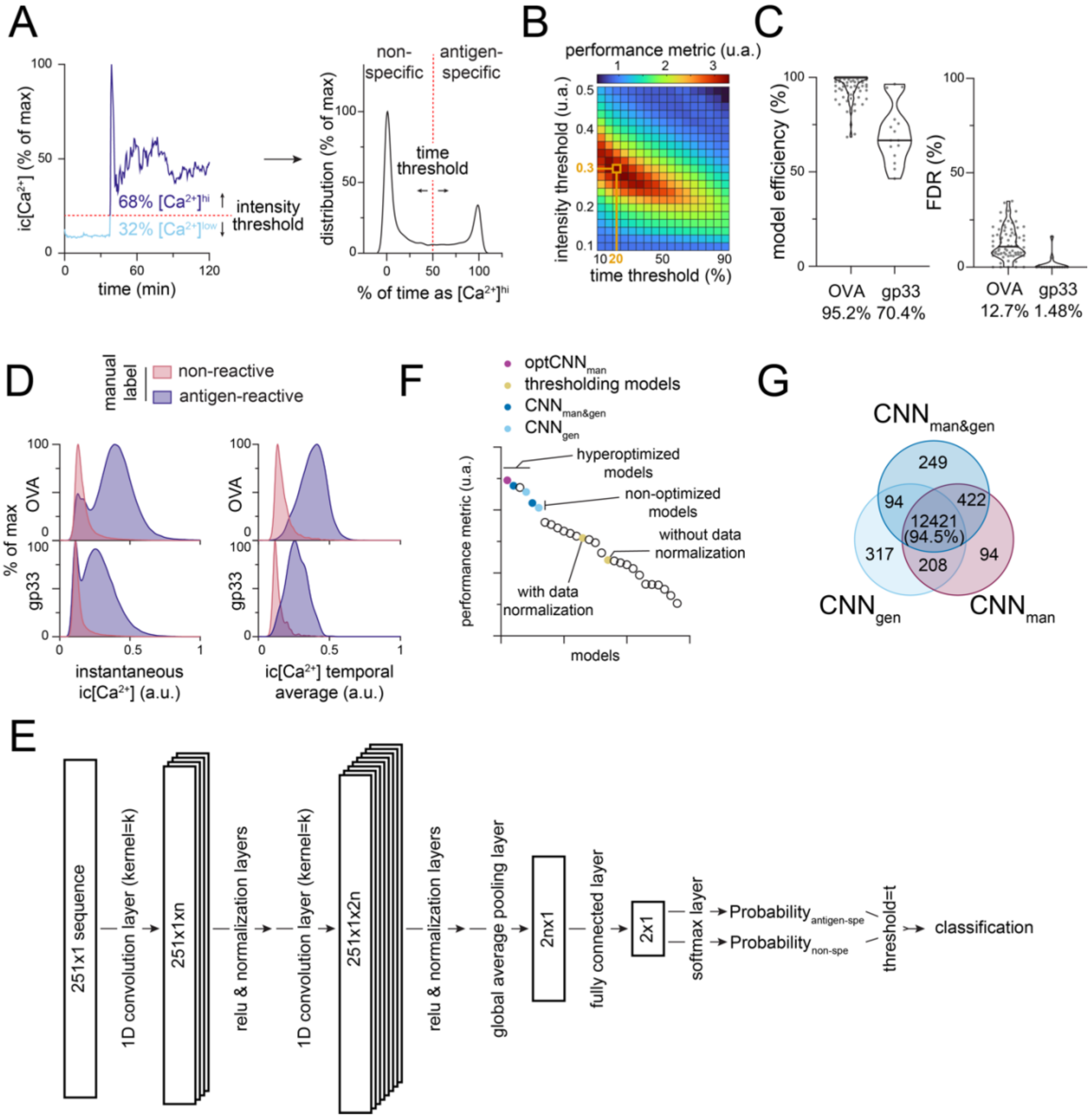


**Fig. S6.** Deep-learning approaches perform better than conventional methods for the classification of T cell activation based on $Ca^{2+}$ fluctuations. (A-C) Use of conventional thresholding methods to classify T cell antigen-reactivity. (A) Varying both intensity and time thresholds (red lines) to predict antigen-specific from non-specific T cells - cells above the time threshold being classified as antigen-specific - we evaluate the performance (on the training dataset) for each pair of thresholds allows to pick an optimal pair of thresholds. (C) Performance of the prediction of antigen-specificity using each pair of thresholds during the training phase. The optimal pair (0.3, 20) of thresholds is highlighted with a thicker line. (C) Using these optimal thresholds, the model efficiency (frequency of cells labeled as antigen-reactive predicted as antigen-specific) and False Discovery Rate (FDR = frequency of cells mispredicted as antigen-specific) is computed on the evaluation dataset. Horizontal lines in the violin plot show the median and numbers below show the average of the distribution. Individual fields of view are represented in gray. (n=73 OVA fields of view; n=15 gp33 fields of view ) (D) Distribution of intracellular $Ca^{2+}$ concentration for all timepoints (left) or the average $Ca^{2+}$ concentration over the whole timelapse (right). according to manual labelling assignment and the peptide used in the co-culture. Each distribution is normalized to its mode. (OVA: n=4584 antigen-reactive cells, n=4864 non-reactive cells; gp33: n=1108 antigen-reactive cells, n=1168 non-reactive cells) (E) Detailed structure of optCNN$_{man}$. Parameters (n, k & t) are indicated in Supp. Table 2. (F) Performance metrics of all models shown in Supp. Table 2 and ranked according to their performance. A few selected models are highlighted. (G) Venn diagram showing the overlap between predictions of the 3 hyperoptimized models: CNN$_{man}$, CNN$_{gen}$ & CNN$_{man\&gen}$, the subscript referring to the ground truth used for training (see Supp. Table 2).

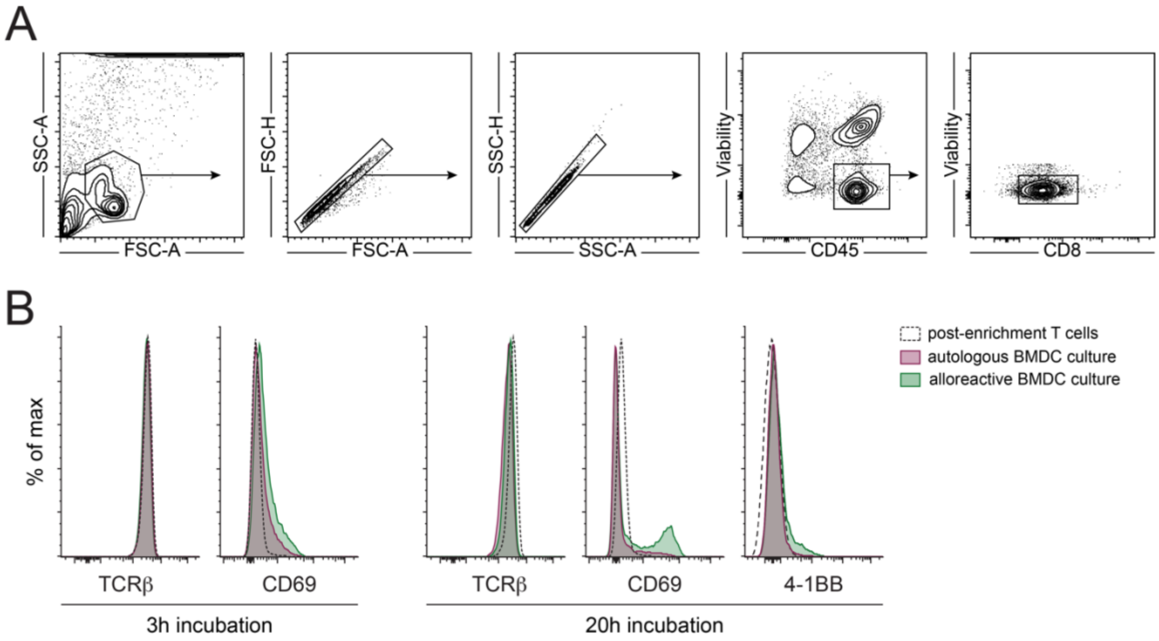


**Fig. S7.** Evaluation of polyclonal T cell activation by flow cytometry. (A) Gating strategy used for the identification of $CD8^+$ T cells following MLR co-culture. (B) Representative distribution of TCRβ, CD69 and 4-1BB expression by T cells in autologous and alloreactive culture conditions, after 3 and 20 hours of co-culture as indicated.

**Table S1.** Distribution of all timelapses into training and evaluation dataset, highlighting the culture peptide, cells stained with CTFR and the number of positive and negative cells in each group.

| | timelapses | | | # cells | |
|---|---|---|---|---|---|
| | peptide | CTFR | # | genotype | manual label |
| training dataset | OVA | P14 | 29 | | |
| | | OT1 | 27 | 3513 antigen-specific | 2673 antigen-reactive |
| | No peptide | P14 | 6 | 6252 non-specific | 7096 non-reactive |
| | | OT1 | 6 | | |
| evaluation dataset | OVA | P14 | 52 | 4584 antigen-specific | 3380 antigen-reactive |
| | | OT1 | 21 | 4864 non-specific | 6068 non-reactive |
| | gp33 | P14 | 6 | 1108 antigen-specific | 594 antigen-reactive |
| | | OT1 | 9 | 1168 non-specific | 1682 non-reactive |

**Table S2.** Training method, performance and ranking based on the weighted performance metric of selected ML models generated during systematic evaluation of ML model structures.

| | | ground truth | hyperoptimization | data normalization | use of derivative | use of speed | data augmentation | efficiency (OVA) | FDR (OVA) | efficiency (gp33) | FDR (gp33) | performance metric (u.a.) | ranking |
|---|---|---|---|---|---|---|---|---|---|---|---|---|---|
| exploration of optimal ML architecture | **clustering approaches** | | | | | | | | | | | | |
| | k-means clustering | gen | | | | | | 49.8 ± 1.87 | 9.78 ± 1.77 | 45.0 ± 4.00 | 3.32 ± 2.67 | 1.283 | 24 |
| | KNN classification | gen | | | | | | 84.4 ± 1.33 | 6.25 ± 0.79 | 42.0 ± 4.94 | 0.75 ± 0.56 | 1.620 | 22 |
| | **SVM based approaches** | | | | | | | | | | | | |
| | SVM classifier | gen | | | | | | 84.4 ± 1.43 | 8.80 ± 0.91 | 44.7 ± 4.85 | 0.74 ± 0.42 | 1.645 | 20 |
| | SVM classifier | gen | Y | | | | | 84.7 ± 1.37 | 8.94 ± 0.91 | 44.2 ± 4.75 | 0.68 ± 0.42 | 1.632 | 21 |
| | SVM classifier | gen | Y | Y | | | | 85.1 ± 1.31 | 8.81 ± 0.95 | 71.3 ± 2.96 | 0.91 ± 0.55 | 2.639 | 14 |
| | recursive SVM classifier | gen | Y | Y | | | | 78.6 ± 1.65 | 7.02 ± 0.81 | 61.6 ± 3.39 | 0.59 ± 0.34 | 2.142 | 18 |
| | recursive SVM classifier | gen | Y | Y | Y | | | 74.8 ± 1.90 | 5.21 ± 0.94 | 67.2 ± 3.14 | 0.92 ± 0.60 | 2.323 | 16 |
| | recursive SVM classifier | man | Y | Y | Y | | | 56.2 ± 2.60 | 3.28 ± 0.70 | 49.0 ± 3.53 | 0.12 ± 0.12 | 1.480 | 23 |
| | **ensemble approaches** | | | | | | | | | | | | |
| | tree classifier | man | Y | Y | Y | | | 87.1 ± 1.47 | 12.6 ± 1.27 | 91.3 ± 1.62 | 18.1 ± 2.09 | 1.935 | 19 |
| | KNN classifier | man | Y | Y | Y | | | 31.6 ± 2.50 | 9.11 ± 2.94 | 35.0 ± 1.93 | 2.96 ± 1.48 | 1.033 | 25 |
| | discriminant classifier | man | Y | Y | Y | | | 87.9 ± 1.24 | 9.82 ± 1.20 | 87.8 ± 2.17 | 14.0 ± 1.88 | 2.383 | 15 |
| | **deep learning approaches** | | | | | | | | | | | | |
| | CNN | man | | Y | | | | 95.2 ± 0.58 | 7.88 ± 0.85 | 90.4 ± 1.58 | 8.27 ± 1.77 | 3.610 | 7 |
| | 1D-Alexnet | man | | Y | | | | 77.1 ± 1.89 | 4.92 ± 1.00 | 63.9 ± 3.63 | 0.92 ± 0.64 | 2.266 | 17 |
| | LSTM | man | | Y | | | | 88.9 ± 1.24 | 5.60 ± 0.84 | 71.9 ± 3.59 | 0.17 ± 0.12 | 3.059 | 12 |
| | CNN-LSTM | man | | Y | | | | 85.2 ± 1.62 | 5.72 ± 1.00 | 72.6 ± 2.99 | 0.52 ± 0.43 | 2.964 | 13 |
| | FC-NN | man | | Y | | | | 88.8 ± 1.27 | 8.82 ± 0.96 | 79.5 ± 2.57 | 1.32 ± 0.55 | 3.258 | 10 |
| exploration of optimal training paramteres | **ground truth exploration** | | | | | | | | | | | | |
| | CNN | man | | Y | | | | 95.2 ± 0.58 | 7.88 ± 0.85 | 90.4 ± 1.58 | 8.27 ± 1.77 | 3.610 | 7 |
| | CNN | gen | | Y | | | | 91.8 ± 1.08 | 6.60 ± 0.78 | 77.8 ± 2.49 | 2.00 ± 0.66 | 3.539 | 8 |
| | recursive CNN | gen | | Y | | | | 90.9 ± 1.11 | 5.03 ± 0.71 | 74.8 ± 2.93 | 1.29 ± 0.65 | 3.430 | 9 |
| | CNN | gen | | Y | Y | | | 94.3 ± 0.76 | 6.27 ± 0.71 | 81.1 ± 2.68 | 1.85 ± 0.77 | 4.072 | 6 |
| | CNN | gen | | Y | | Y | | 91.4 ± 1.07 | 8.35 ± 0.78 | 77.0 ± 2.78 | 2.82 ± 0.71 | 3.175 | 11 |
| | CNN | man&gen | | Y | | | | 91.5 ± 1.07 | 4.82 ± 0.74 | 81.0 ± 2.48 | 1.57 ± 0.91 | 4.226 | 5 |
| | **hyperoptimization** | | | | | | | | | | | | |
| | **CNN** | **man** | **Y** | **Y** | | | | **93.7 ± 0.83** | **6.26 ± 0.82** | **88.6 ± 1.97** | **2.90 ± 1.34** | **4.944** | **1** |
| | CNN | gen | Y | Y | | | | 95.1 ± 0.72 | 6.75 ± 0.75 | 86.2 ± 1.85 | 2.64 ± 0.76 | 4.577 | 4 |
| | CNN | man&gen | Y | Y | | | | 91.4 ± 1.09 | 5.73 ± 0.84 | 87.2 ± 1.57 | 2.72 ± 1.50 | 4.773 | 2 |
| | **data augmentation** | | | | | | | | | | | | |
| | CNN | man | Y | Y | | | Y | 93.1 ± 0.90 | 6.04 ± 0.80 | 86.3 ± 2.17 | 2.52 ± 1.02 | 4.717 | 3 |

First, we explore amongst well known ML architectures the best performing model with our dataset. We use the term recursive to describe the process of training a first model on $Ca^{2+}$ fluctuation, the prediction of which is used as ground-truth for a second SVM model. (‘man’ = manual label used as ground-truth; ‘gen’= genotype used as ground-truth; ‘man&gen’ = antigen-specific T cells manually labeled as antigen-reactive used as ground-truth) Data shows average ± SEM. (KNN = k-nearest neighbors, SVM = Support Vector Machine, LSTM = Long Short Term Memory networks, FC-NN = fully connected neural networks, CNN = Convolutional Neural Network).

Table S3. Parameters used for the generation, the training, and the prediction of antigen-specificity with the three hyperoptimized models described in the study.

| ML structure | man | gen | man&gen |
|---|---|---|---|
| number of neurons (n) | 30 | 30 | 60 |
| kernel size (k) | 8 | 8 | 2 |
| number of classes | 2 | 2 | 3 |
| **training parameters** | | | |
| optimizer | adam | adam | adam |
| minibatch size | 100 | 25 | 50 |
| internal validation | 15% | 15% | 15% |
| **prediction** | | | |
| threshold (t) | 0.5 | 0.5 | 0.4 |

Letters in parentheses reflect parameters presented in Supp. Fig. 6
